# Systematic identification of novel regulatory interactions controlling biofilm formation in the bacterium *Escherichia coli*

**DOI:** 10.1101/155432

**Authors:** Gerardo Ruiz Amores, Aitor de las Heras, Ananda Sanches-Medeiros, Alistair Elfick, Rafael Silva-Rocha

## Abstract

Here, we investigated novel interactions of three global regulators of the network that controls biofilm formation in the model bacterium *Escherichia coli* using computational network analysis, an *in vivo* reporter assay and physiological validation experiments. We were able to map critical nodes that govern planktonic to biofilm transition and identify 8 new regulatory interactions for CRP, IHF or Fis responsible for the control of the promoters of *rpoS*, *rpoE*, *flhD*, *fliA*, *csgD* and *yeaJ*. Additionally, an *in vivo* promoter reporter assay and motility analysis revealed a key role for IHF as a repressor of cell motility through the control of FliA sigma factor expression. This investigation of first stage and mature biofilm formation indicates that biofilm structure is strongly affected by IHF and Fis, while CRP seems to provide a fine-tuning mechanism. Taken together, the analysis presented here shows the utility of combining computational and experimental approaches to generate a deeper understanding of the biofilm formation process in bacteria.

## INTRODUCTION

Bacteria shift from their free-swimming lifestyle to adopt a community structure to benefit from the micro-environment created in a biofilm ^1^, gaining protection against hazardous substances, and leading in some cases, to antibiotic resistance ^1, 2, 3^. It is well-understood that when differentiating from the planktonic to a biofilm structure bacteria transit through specific stages, with each functional stage accompanied by changes to expression of specific genes of the flagella-biofilm regulatory network ^1,2,3^. In *Escherichia coli*, the complex transcriptional regulatory network of flagella function and curli fimbriae production, the principal biofilm structure indicator, has been investigated in various reports ^4, 5, 6,7^. In this organism, the process is controlled by the RpoS sigma factor and FlhDC regulator. These master regulators receive major regulatory inputs from c-di-GMP, cAMP and ppGpp, which are modulated by a series of environmental and physiological stresses ^8,9,10^. In this sense, the *flhDC* genes are expressed at post-exponential phase and their products control more than 60 genes involved in flagella synthesis and related functions, such as chemotaxis ^11^. Also at stationary phase, the general stress response master regulator RpoS is produced and controls over 500 genes in a highly complex regulatory network ^7^. Therefore, the interplay between those master regulators and downstream-activated genes modulates the complex transition between planktonic and biofilm stages.

In the case of planktonic bacteria, mainly found in the exponential phase, flagella maintenance is due to the activation of effector molecules such as FliA sigma factor. Whilst, at stationary phase, biofilm formation is initiated by increasing levels in the secondary messenger c-di-GMP, as well as expression and activation of *rpoS,* which in turn leads to the activation of *csgD* ^5, 12^. Additionally, there are alternative mechanisms that involve different key genes of the biofilm pathway, also leading to *csgD* activation and scale-up of curli fiber production; such as CpxR and ClpX which play a complex dual role during bacterium development to inhibit or activate both programs ^8, 10, 13^. Undoubtedly, the interplay between these molecules acts to modulate the proper execution of the flagella-biofilm program^14, 15^. Experimentally, the transition between flagella and biofilm formation is evaluated using a “batch cells” assay, in which cells grow and attach to the chemically-inert surface of a microliter dish under static conditions, generating a biofilm in an “aquatic environment”. Additionally, the “macro-colonies” assay allows a bacterial population to grow over extended periods of time on agar plates, leading to striking morphological patterns, resembling biofilms growing in biological materials.

While hundreds of reports have dissected the molecular mechanisms leading to biofilm formation in several bacterial groups, a full understanding the dynamics of the complex regulatory network controlling planktonic to biofilm transition still lies out of reach. This is particularly true since most studies analyzed the effect of particular regulators on the expression of specific sets of genes, without addressing the complexity of the network using a more systematic approach. In particular, since a few global regulators (GRs) are able to control most of the genes in genome of *E. coli*^16^, it is anticipated that the role of these transcriptional factors in the biofilm formation network is rather underestimated. GRs are expressed or activated differentially during growth or under specific input signals ^17^. Specifically, CRP is related to the control of metabolic processes in bacteria and it is differentially activated by substrates ^18^, while IHF (growth rate dependent) and Fis (expressed at early exponential phase) are dual nucleoid associated proteins (NAPs) that can work both as activators and repressors ^19, 20^. During the transition from motile cells to biofilm structure, a mixture of different developmental stages occur. In this sense, small molecules and GRs direct the activation of signaling proteins that are sensed by neighboring cells, coordinating the construction of this complex community structure ^14, 15^. Therefore, mapping hidden interactions in the flagella-biofilm regulatory network would provide a better understanding of this developmental-like program and provide new targets for intervention and/or engineering ^21, 22^.

In this work, we used computational tools and molecular biology, together with microbiology approaches to investigate the effect of the GRs CRP, IHF and Fis on the *E. coli* flagella-biofilm network. First, to analyze regulatory interactions already reported, we reconstructed the flagella-biofilm transcriptional regulatory network *in silico* using previous data and we analyzed its structure. Next, by using promoter expression assays, we mapped novel regulatory interactions between the three GRs and key genes controlling biofilm formation. Finally, novel regulatory interactions mapped using this approach were added to the original network to generate an enhanced appreciation of biofilm formation. The general strategy used here is depicted in **Fig. 1**. These results suggest that IHF and Fis are important modulators of the process while CRP seems to exert a fine-tuning effect. Additionally, using motility, adherence and biofilm formation assays, we validated key regulatory effects observed using transcriptional fusions. All together, the systematic investigation presented here adds new understanding of the complex regulatory program that controls biofilm formation in *E. coli*, revealing a novel player in this network.

**Figure 1.**
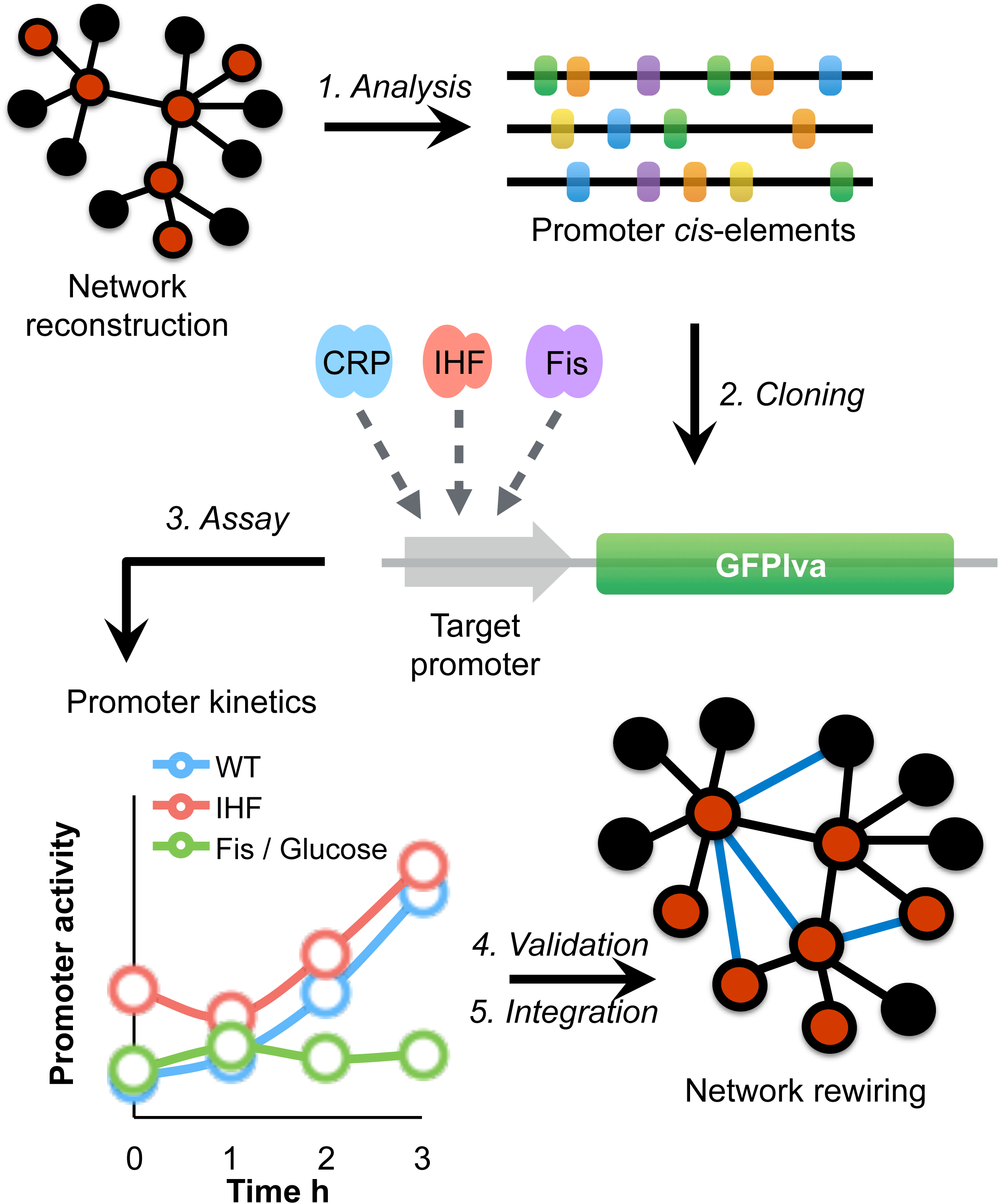
General strategy to define novel regulatory interactions controlling planktonic/biofilm transition. The flagella-biofilm network was constructed and analyzed based on available data from the literature. Subsequently, analysis of the promoter architecture regions of the principal nodes effectors was performed to confirm the interactions reported. Next, cloning of the natural promoter regions in the pMR1-reporter system were developed. Promoter activity was determined for each promoter in different conditions as established in material and methods. Finally, GFP activity was transform into connectivity data and loaded into flagella-biofilm network to gain a better understanding of this program.

## RESULTS AND DISCUSSION

### *In silico* analysis of the regulatory network controlling planktonic to biofilm transition in *E. coli*

To reconstruction the flagella-biofilm regulatory network, information regarding the transcriptional regulation of *E. coli* was gathered from scientific articles and compiled with that from RegulonDB ^23^ and KEGG DB ^24^. The resulting network consisted of 37 nodes representing the connections between different transcription factors (TFs), GRs and small molecules **(Fig. 2)**. The degree analysis (which shows the most connected nodes) of the flagella-biofilm transcriptional regulatory network identified three major hubs, *rpoS*, *flhD* and *csgD* (**Fig. S1A**). The betweenness and edge betweenness analysis which provides information about the most important nodes and paths in the network, shows that from 37 nodes, 8 seem to be the principal effectors that modulate the proper gene expression of the network (*rpoS, csgD,* c*-*di*-*GMP, *rpoE, cpxR, clpX, flhD, fliA*). Interestingly, *rpoS* is the main effector in the network and it is modulated by different inputs. From those, c-di-GMP presents high betweenness and edge connection values (**Fig. 2**). From the 37 nodes, *rpoD* and CRP were the main nodes, which exert more out-regulatory signals to other nodes, while all others possessed fewer out-connections (**Fig. S1B**). Additionally, this analysis shows three paths that seem to be important to maintain balance in the network. The first critical connection that maintains flagella-biofilm program equilibrium lies between *rpoS* and *rpoE*, activating both *fliA* for motility and its repressor *matA*, to eventually develop adherence. Secondly, *rpoS* with the TF *cpxR*, which modulates *rpoE*, but also has an important connection with the TF *nsrR*, which represses *fliA* expression. Thirdly, the activation of *clpX* ATPase by *rpoS* that exerts *flhD* inhibition at post-transcriptional and/or post-translational levels ^25^. From the analysis presented here, we selected ten genes in order to perform a systematic investigation of the effect of CRP, IHF and Fis, which are among the main GRs of the regulatory network of *E. coli* ^26^. We expected that this approach would reveal some hidden interactions controlling the critical nodes of the flagella-biofilm network. The selected genes of interest were *rpoS, rpoE, csgD, cpxR, flhD, fliA, matA, ompR, adrA and yeaJ*. In order to analyze the promoter activity of the selected genes, the promoter regions of the genes were cloned upstream of a GFPlva reporter system ^27^ and used to transform wild-type and *Δihf* and *Δfis* mutant strains of *E. coli*. As control, an empty reporter plasmid and a strong synthetic promoter named BBa_J23100 (http://parts.igem.org/Part:BBa_J23100, here referred as *Pj100*) were also used to transform the same strains (**Fig. S2**). The 36 reporter strains were then analyzed as presented in the next section.

**Figure 2.**
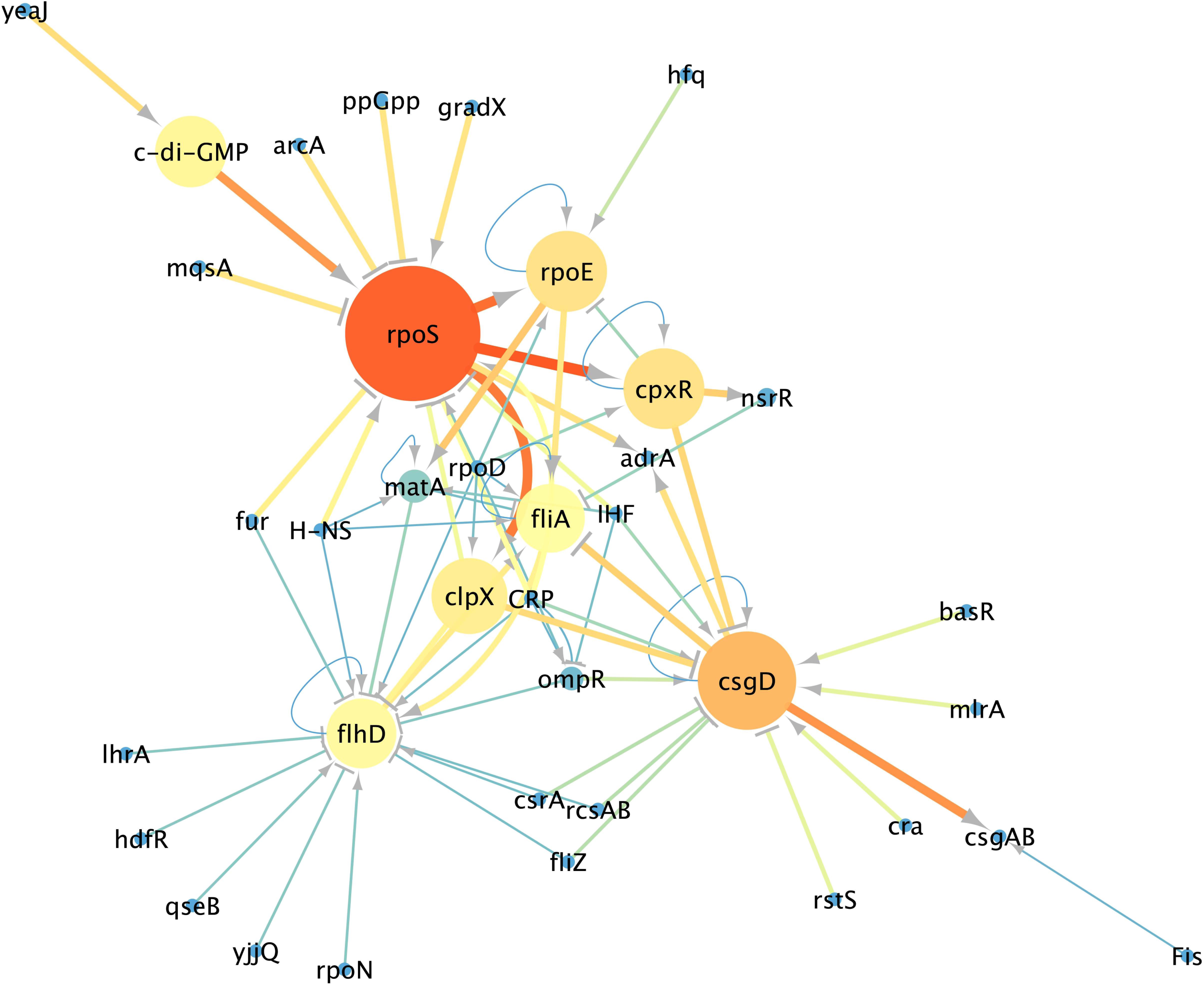
Flagella-biofilm transcriptional regulatory network. The principal nodes and paths, which drive the flagella-biofilm network were analyzed using Cytoscape 3.4.1. The network was analyzed by using betweenness centrality and edge betweenness centrality algorithms ^54^. Size of the nodes (circles) indicates betweenness analysis and width of the Edges (lines connections the circles) indicates the edge betweenness analysis. Scale colors from red to bright to dark indicate the high to low values.

### CRP, IHF and Fis modulate promoters of *rpoS*, *rpoE* and *flhD* master regulators

We started our investigation by considering the three putative master regulators of the transition from planktonic to biofilm phases. The sigma factor RpoS has been proposed as a main regulator of the flagella-biofilm network ^7,8^, in agreement with our network analysis. Thus, we were interested to understand the effect of these three GRs on the promoter region of the *rpoS* gene. For this, overnight cultures of *E. coli* wild-type strain carrying the transcriptional *PrpoS::GFPlva* fusion were diluted 1:100 in fresh M9 minimal medium supplemented with glycerol as sole carbon source, and GFP fluorescence was measured at 20 minute time intervals over 8 hours at 37°C. As can be seen in **Fig. 3A**, the analyzed promoter presented a growth-phase-dependent activity with increased activity toward the end of the growth curve. When the same reporter system was analyzed in *E. coli Δihf* and *Δfis* mutant strains, we observed a significant increase in promoter activity in the absence of IHF and a strong activity in the absence of Fis (**Fig. 3A left panel**, red and green lines, respectively). These data strongly suggest that IHF and Fis GRs are acting to repress the *rpoS* promoter. To understand the role of CRP over the *rpoS* promoter region, we employed the glucose inducible CCR system to down-modulate the activity of CRP by adding glucose in the same experimental conditions as described above ^28^. When the three reporter strains were analyzed in the presence of 0.4% glucose, we observed the same general expression profile as in **Fig. 3A left panel** but a remarkably reduced promoter activity (**Fig. 3A right panel**), confirming the previously reported positive role of CRP on the *rpoS* promoter ^29^. Additionally, since the synthetic constitutive promoter (*Pj100*) was not significantly affected in the strains and conditions used (**Fig. S2**), we concluded that the effects observed for *rpoS* promoter are true regulatory events taking place at this element.

**Figure 3.**
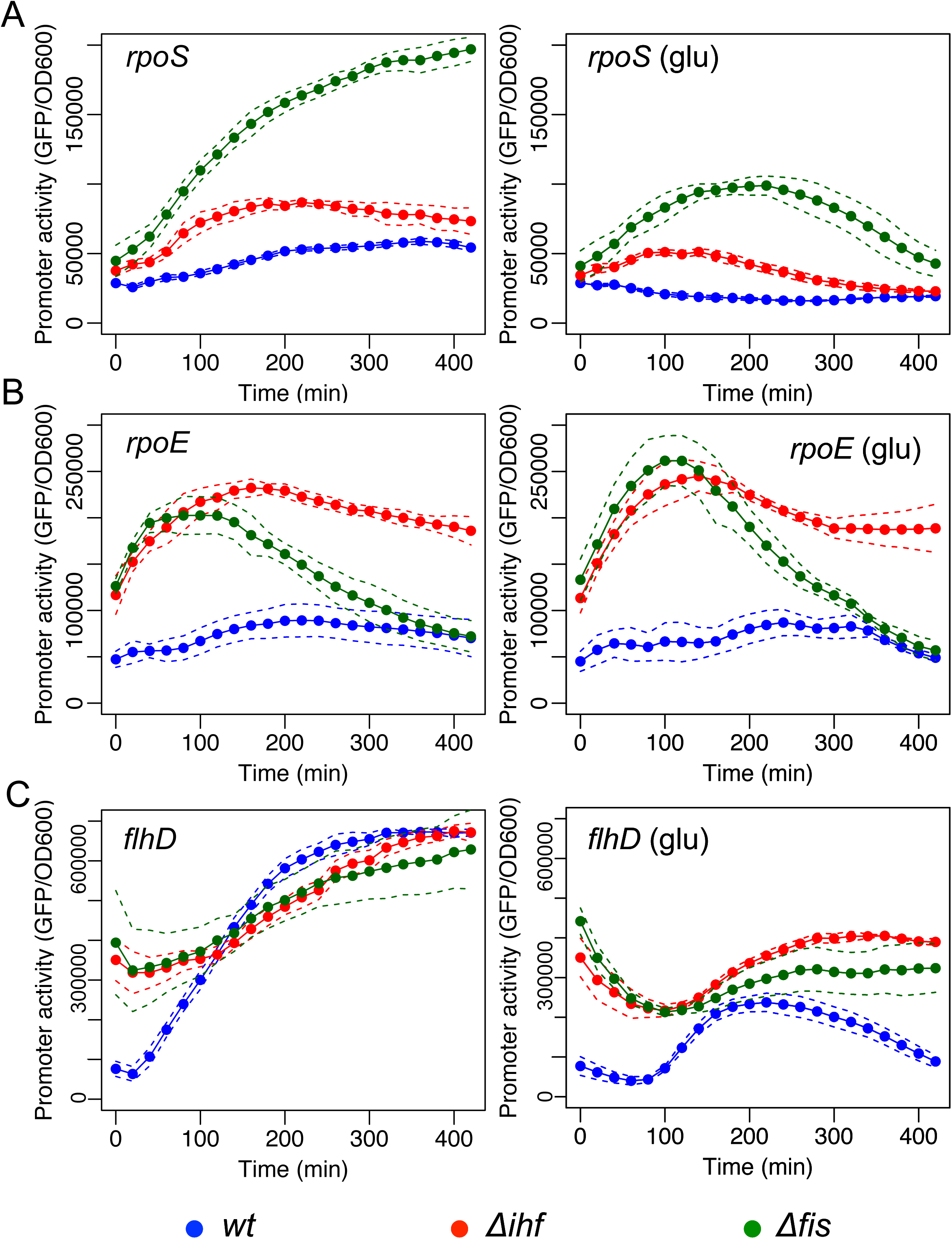
Effect of CRP, IHF and Fis GRs over the promoter activity of *rpoS*, *rpoE* and *flhD*. Promoter activity assay of (A) pMR1-*PrpoS,* (B) pMR1-*PrpoE,* and (C) pMR1-*PflhD* were evaluated in *E. coli* BW25113 *wild type* (blue line), Δ*ihf* (red line) and Δ*fis* (green line) in 96well plate as described in methods in the absence (left panel) or presence (right panel) of 0.4% of glucose. GFP fluorescence was measured each 20 minutes at 37 °C during 8 hours in static conditions and normalized by OD600. Solid lines represents the average from three independent experiments while dashed lines are the upper and lower limits of standard error of the mean (S.E.M).

When we analyzed the activity of the *rpoE* promoter, which controls the expression of a sigma factor linking RpoS activity with flagella genes (**Fig. 2)**, we observed a strong increase in promoter activity in the *Δihf* and *Δfis* strains of *E. coli* when compared to the wild-type strain (**Fig. 3B left panel**), indicating that IHF and Fis have a negative effect on the activity of this promoter. However, contrary to the observation for *rpoS*, the addition of glucose (**Fig. 3b right panel**) to the media (which inhibits CRP activity) did not result in any significant change in *rpoE* promoter activity.

After investigating the activity of *rpoS* and *rpoE* promoters, we analyzed the promoter of *flhD* gene, which encodes a master regulator of flagellar genes and of the flagella-related *fliA* sigma factor ^30^. Using the same experimental conditions as described above, we first observed that *flhD* promoter activity has a strong growth-dependence, with activity increasing during the growth curve when assayed in the wild-type strain of *E. coli* (**Fig. 3C left panel**). When, we assayed the promoter activity in *Δihf* and *Δfis* mutant strains, we observed a similar expression profile as in the wild-type, with the exception that in both strains the initial promoter activity was significantly higher than in the wild-type. When we assayed promoter response in the three strains in the presence of glucose (to trigger CRP inactivation), activity was strongly impaired in all strains (**Fig. 3C right panel**), confirming the reported role of CRP as activator of *flhD* expression ^31^.

### Novel regulatory effects for CRP, IHF and Fis exerted on *csgD*, *fliA* and *yeaJ* promoters

Once we had investigated the effect of the GRs of flagella to biofilm transition, we decided to analyze the promoters of genes related to biofilm formation and an additional lower flagella regulator (*fliA*). CsgD is a central protein related to, and an indicator of, biofilm formation ^12,32^. When we analyzed the activity of *csgD* promoter in the wild-type strain, we observed a peak of expression at about 200 min of growth with a reduced activity later in the growth phase (**Fig. 4A left panel**). Investigating this promoter in *Δihf* and *Δfis* mutants, we observed a strong decay in promoter activity, indicating that both proteins positively modulate the activity of the promoter. Moreover, when we added glucose to the growth media (**Fig. 4A right panel**), we observed a generalized decay in promoter activity in all three strains, indicating a positive role of CRP for this promoter. Reflecting on these results, it is interesting to note that, while CRP and IHF have been reported as positive regulators of the *csgD* promoter ^33,34^, the positive effect of Fis has never been demonstrated before.

**Figure 4.**
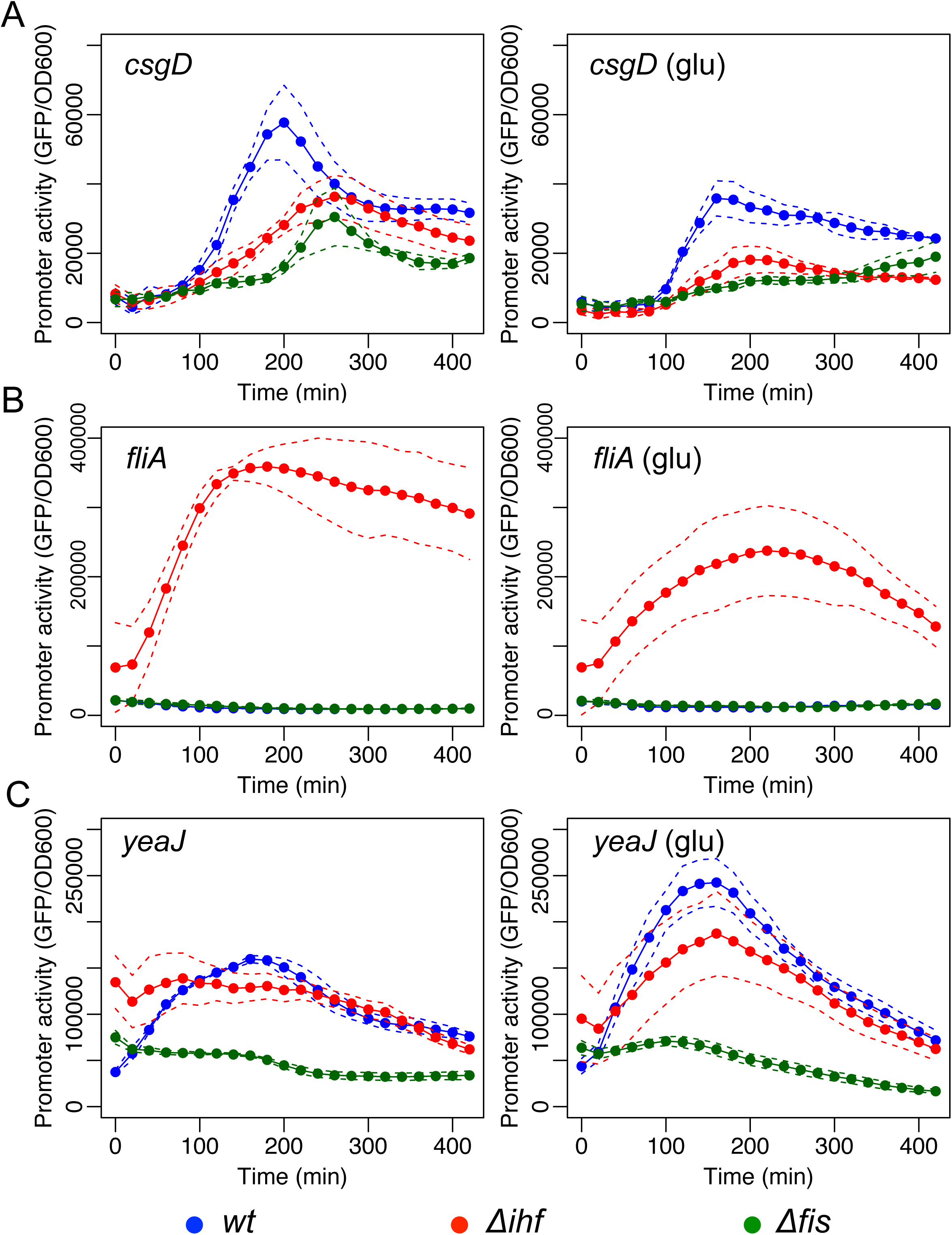
Effect of CRP, IHF and Fis GRs on the promoter activity of *csgD*, *fliA* and *yeaJ*. GFP promoter activity assay of (A) pMR1-*PcsgD,* (B) pMR1-*PfliA,* and (C) pMR1-*PyeaJ* were evaluated in *E. coli* BW25113 *wt* (blue line), Δ*ihf* (red line) and Δ*fis* (green line) in 96well plate as described in methods in the absence (left panel) or presence (right panel) of 0.4% of glucose. GFP fluorescence was measured each 20 minutes at 37 °C during 8 hours in static conditions and normalized by OD600. Solid lines represents the average from three independent experiments while dashed lines are the upper and lower limits of standard error of the mean (S.E.M).

In the case of the *fliA* gene that codes a sigma factor specifically related to flagella genes, analysis of promoter activity in both wild-type and *Δfis* mutant strains revealed a very low level of activity throughout the growth curve, both in the presence and absence of glucose (**Fig. 4B**). However, in the *Δihf* mutant strain, this promoter displays a strong increase (30-fold) in activity in the absence of glucose (left panel), while this level was lower when glucose was added (right panel). Since *fliA* expression is dependent on FlhD ^30^, the observed decay of *fliA* promoter activity during growth in the presence of glucose could be the result of a cascade process, with the apparent IHF repression of this promoter suggesting a previously unreported regulatory interaction. To test the direct effect of IHF on FliA, we identified two putative binding sites for this global regulator at *fliA* promoter **Fig. S3A**. We then constructed a mutant version of this promoter with 11 point mutations that fully abolish both sites and tested this new variant as before. As shown in **Fig. S3B**, expression of the mutated version of *fliA* promoter was low in the wild-type and *Δfis* mutant strains at the same level as the original promoter, and displayed a higher activity in the *Δihf* mutant, both in the absence and presence of glucose. These data suggest that the effect of IHF on *fliA* promoter is indirect.

We next analyzed the effect of GRs on the expression of *yeaJ*, a diguanylate cyclase coding gene, which modulates the levels of c-di-GMP in the cell. In *E. coli*, deletion of *yeaJ* results in strains with reduced motility at 37 °C ^8^, however no regulatory proteins have been demonstrated to play a role in the expression of this gene. The expression profile of the *yeaJ* promoter in wild-type and Δ*ihf* mutants strains showed a strong peak of activity at 200 min of experiment (**Fig. 4C**). However, when this promoter was assayed in the Δ*fis* mutant, we observed a significant decrease in promoter activity, indicating that the Fis regulator could play a positive role in its expression. Furthermore, addition of glucose to the growth media (**Fig. 4C right panel**) resulted in an increase in the promoter activity in the first minutes of growth, suggesting the existence of additional, potentially CRP-dependent, regulatory mechanisms at this promoter. Finally, the additional three promoters selected for investigation (from *adrA*, *cpxR* and *ompR* genes, **Fig. S4**) displayed maximal activity very similar to that of the negative control (the empty reporter vector, pMR1 in **Fig. S2**), which precludes unequivocal conclusions concerning their regulation.

### Reconstruction and analysis of the regulatory network with novel interactions

In order to generate fresh insight into the regulatory network controlling biofilm formation in *E. coli*, we added the new regulatory interactions identified in the previous sections to the network from **Fig. 2.** For this, the effect of CRP, IHF and Fis over the promoter regions obtained by our promoter activity assay was transformed into activation or inhibition links. From the 27 interactions tested, we suggested 8 new interactions, and were able to confirm 5 described interactions (**Table S1**). We integrated these new interactions into the network and re-performed the centrality measurements. Using this approach, we observed that the integration of the experimental data changed the topology of the network, while maintaining the three major hubs identified (**Fig. S5A**). Yet, the out-degree analysis showed that *rpoD,* CRP, IHF, and Fis in minor degree are the main nodes exerting the most out-connections to all other nodes (**Fig. S5B**), while the edge betweeness analysis indicates that, upon the addition of the new interactions, the critical nodes of the network remain the same (**Fig. 5**). Taken together, these data suggest the existence of previously unidentified regulatory interactions in the biofilm regulatory network that play a critical role in the planktonic to biofilm transition.

**Figure 5.**
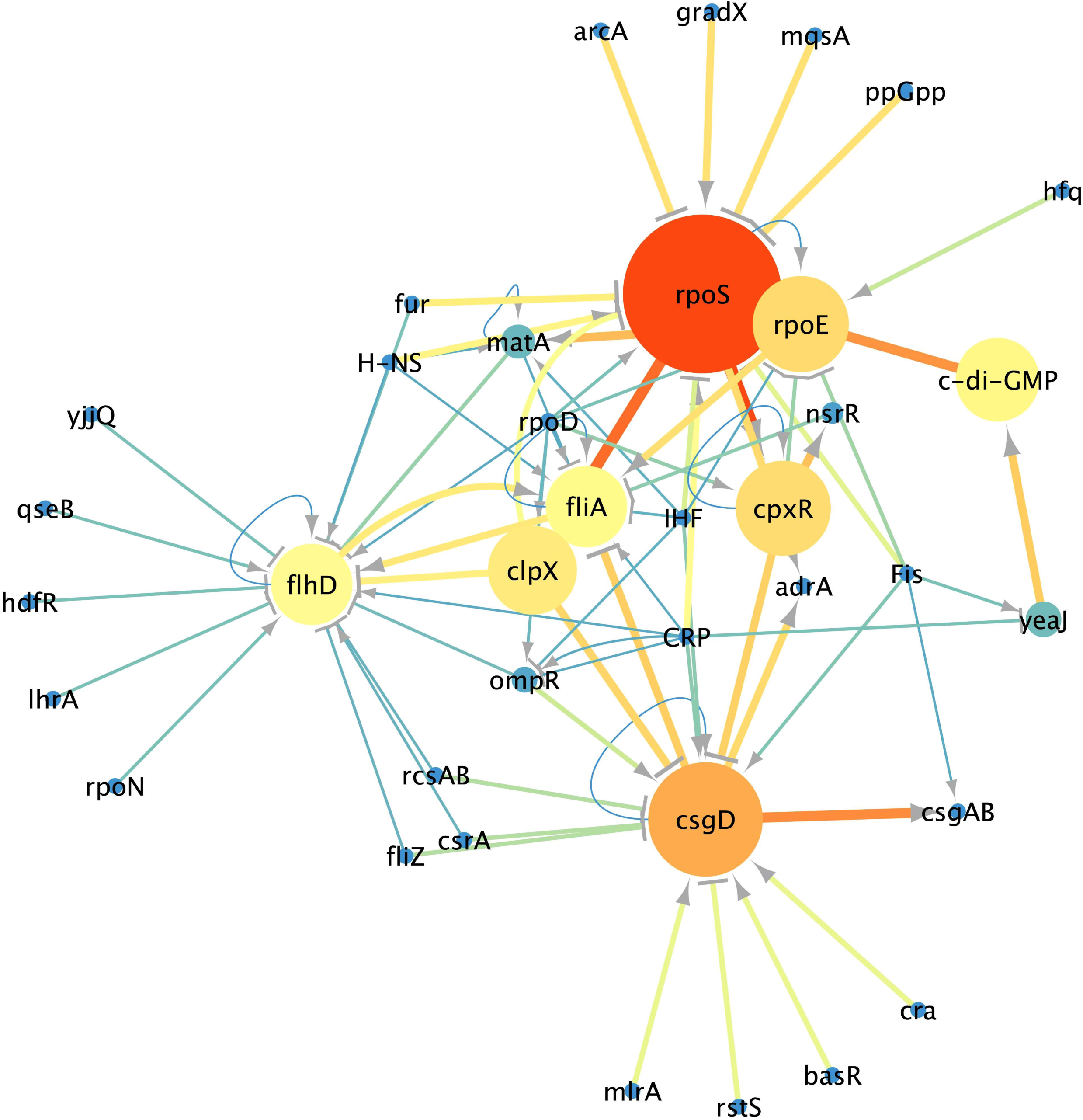
Experimental data integration to the Flagella-biofilm transcriptional regulatory network. Promoter activity values were transformed into activation or repression connections and were loaded into Cytoscape 3.4.1. An organic algorithm was used to the properly cluster visualization. The network was analyzed by use the betweenness centrality and edge betweenness centrality measurements. Size of the nodes (circles) indicates betweenness analysis and width of the Edges (lines connections the circles) indicates the edge betweenness analysis. Scale colors from red to bright to dark indicate the high to low values.

### Phenotypic consequences of loss of GRs for the planktonic and biofilm stages of *E. coli*

Flagella play an important function during the different developmental phases of biofilm formation such as attachment at a surface, as well as bacterial cell cohesion within the biofilm ^2, 30^. Therefore, we were interested in understanding the effect of the GRs CRP, IHF and Fis on the flagella function. As it can be seen in **Fig. 6**, after 24h of incubation in a low agar plate, *E. coli* BW25113 wild-type strain showed reduced motility in the absence of glucose. However, analysis of the Δ*ihf* strain in similar conditions revealed a strong motile phenotype, with this effect more apparent after 24h of incubation with full spreading over the plate observed (**Fig. S6**). Finally, under similar conditions, the Δ*fis* strain presented reduced motility, which is more evident at 24h. It is important to notice that addition of glucose to the media generated a general decrease in motility of all strains (**Figs. 6** and **S6**, right panels). Interestingly, the observed strong phenotypic effect of Δ*ihf* on the motility of *E. coli* can be traced directly to the strong increase of *fliA* promoter activity in this mutant strain, since FliA sigma factor controls several genes related to flagella formation in this organism ^35^.

**Figure 6.**
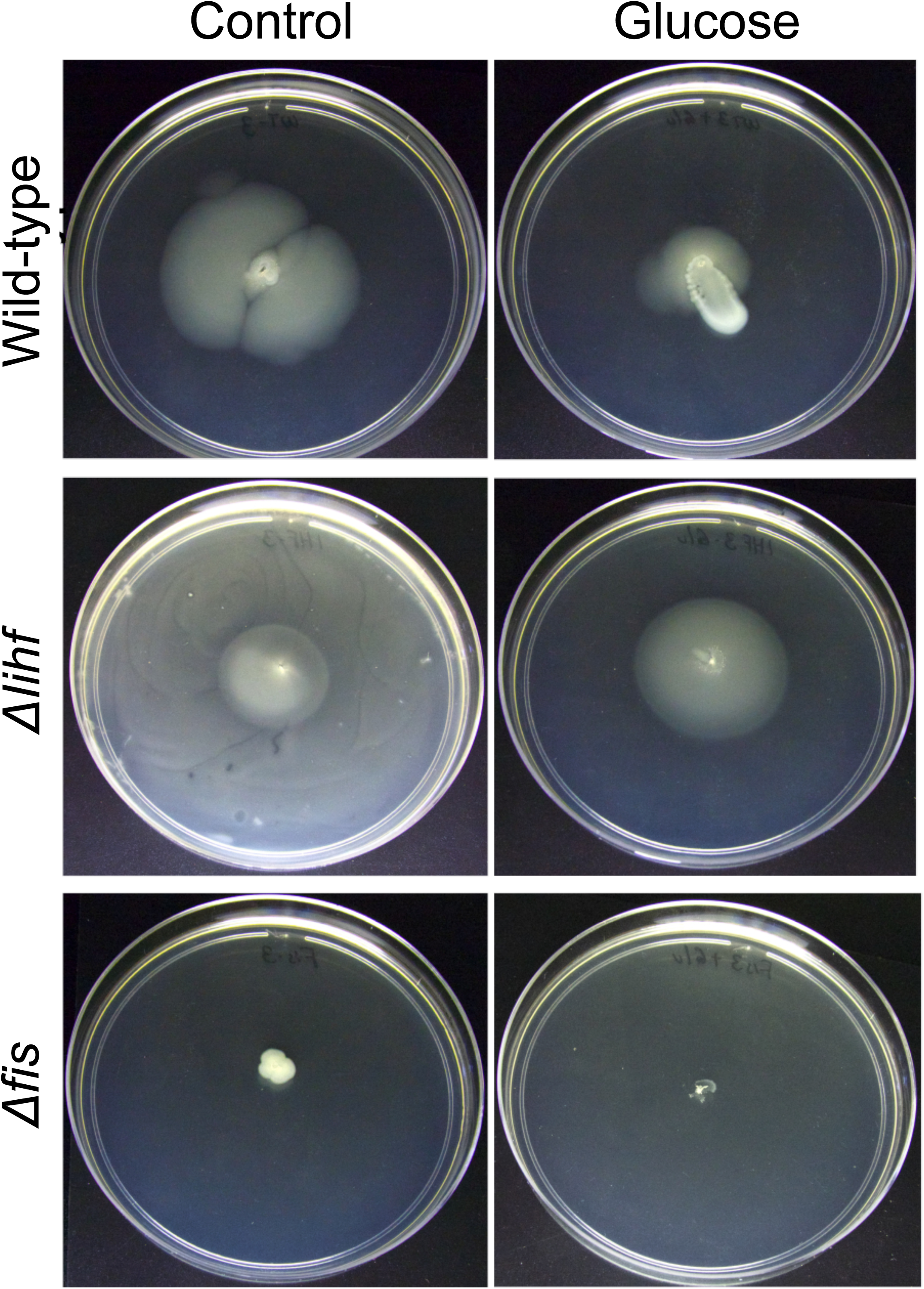
Effect of GRs in the motility program at 24h. Motility phenotype of *E. coli* BW25113 wild-type and mutant strains were evaluated by cell motility assay at 24h in the presence or absence of glucose as depicted. Divergent motility capability is observed between the different conditions, proving the effect of the GRs CRP, IHF and Cis to modulate the motility program. The results are representative of 3 independent experiments.

We next analyzed the capability of the different strains to attach to a solid surface, which is indicative of early biofilm formation, using the protocol described by O’Toole *et al*., 2011. For this, we performed the experiments at 30 °C and 37°C, since different reports have used different temperatures^36, 37^. In general, *E. coli* cells adhered to the wall of the 96-well plates in all conditions. At 37°C, two- or three-fold increases in biofilm yields were observed when compared with *E. coli* cells cultured at 30°C, as can be seen by comparing the biofilm indexes in **Fig. 7A**. In general terms, the *Δihf* mutant strain of *E. coli* displayed decreased adhesion at both temperatures, which is in agreement with the increased motility observed for this strain in **Fig. 6**. By the same token, adhesion of the Δ*fis* mutant strain at 37 °C was significantly increased when compared to the wild-type, which could be due to the reduced motility observed before. In most of the cases, the general adhesion observed for the strains was decreased in the presence of glucose, with the exception of the adhesion level of Δ*ihf* mutant at 30°C. These data suggest that, at 37°C, CRP and IHF are important effectors to activate the adherence program. They also suggest that Fis is acting as a repressor of the early biofilm program, since in its absence, motility is higher at 37 °C. Altogether, adherence yields are better when induced at 37°C than at 30°C, and the three GRs CRP, IHF and Fis participate differentially to regulate the adherence capabilities of *E. coli*.

**Figure 7.**
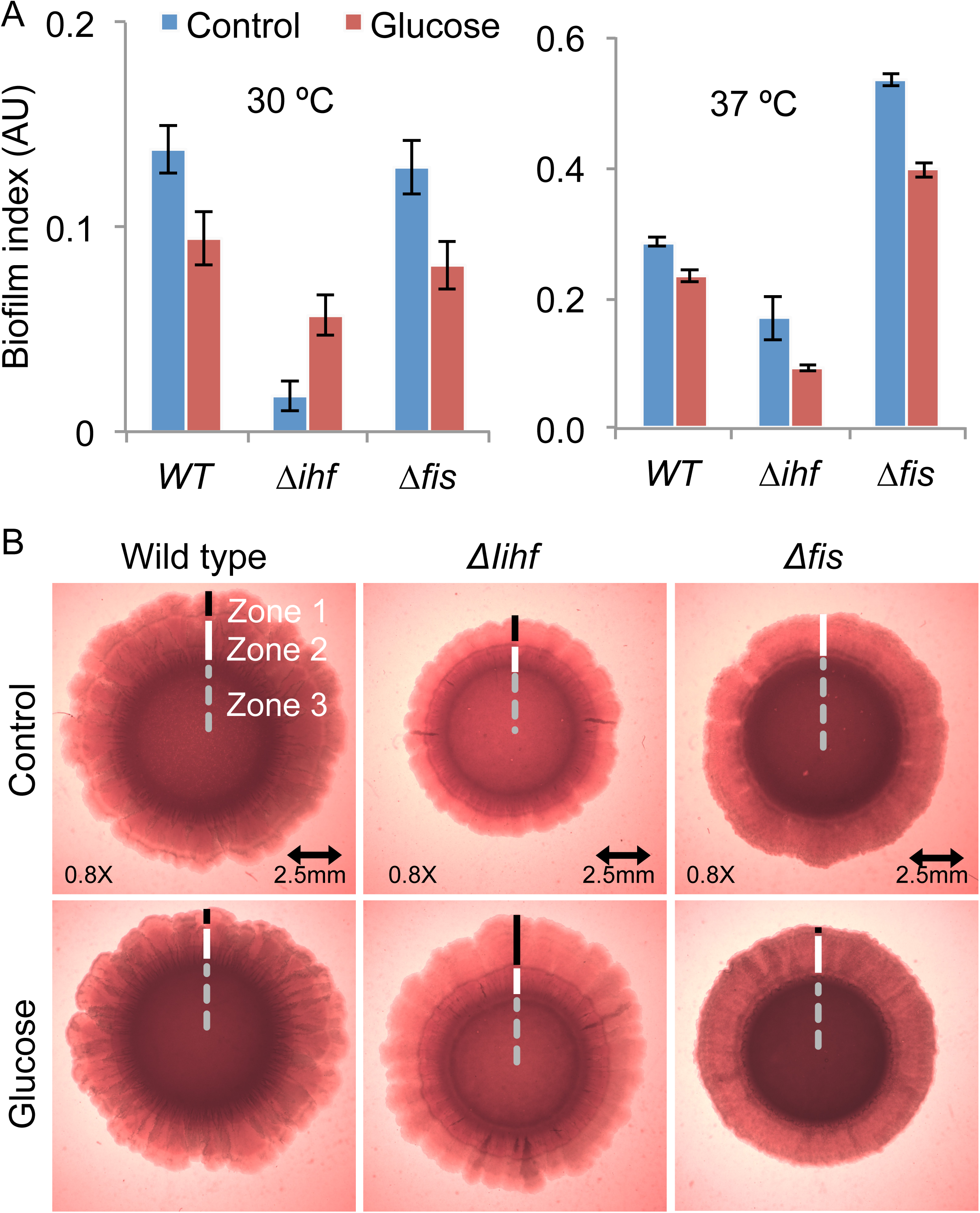
Capability of *E. coli* and mutant strains to develop adherence and mature biofilm. A) Adherence capability of E. coli BW25113 *wild-type*, Δ*ihf* and Δ*fis* were evaluated in 96-well plate using violet crystal method. Comparisons of adherence capability of BW25113 wt and mutant strains at 30 and 37°C are shown. Vertical bars are standard deviations calculated from three independent experiments. B) Mature biofilm formation of *E. coli* BW25113 *wt*, Δ*ihf* and Δ*fis* strains were perform using congo red plate assay. Comparisons of the mature biofilm morphological characteristics of wild type and mutant strains at 37°C in the presence or absence of glucose is shown. Dashed line, Zone III; white line, zone II; black line, zone I; arrows represent wrinkles and clefts structures. The results are representative of 3 independent experiments.

As depicted in **Fig. S7**, all the strains were capable of developing biofilm structures at 30 °C and 37 °C in the presence or absence of glucose, as previously reported. In general, *E. coli* cells grown at 30 °C produced a biofilm structure resembling that reported for 28 °C, while those grown at 37 °C conditions showed a more visually complex biofilm structure with three morphologically distinct zones, as reported by Serra *et al*., 2013 (**Fig. 7B**). More explicitly, in wild-type cells a concentric ring delineates zone 3, which presents an intense red color suggesting curli formation by the different layers of stationary phase cells. Zone 2 represents the intrinsic capability of wild-type community to produce wrinkles and curli. Finally, zone 1 presents weak red color, an indicative of bacteria at exponential phase related with colony expansion but not with curli production ^7^. In the presence of glucose, the biofilm structure formed by the wild-type strain presented slight differences. When we analyzed the *Δihf* mutant strain, we observed a significant reduction in the colony size and degree of pigmentation (that could be related to lower curli production) when compared to the two other strains (**Fig. 7B**). This size reduction can be evidenced by the systematic decrease in the three zones of the colony in this strain. Yet, analysis of biofilm formation in this strain in the presence of glucose showed a recovery in colony size generated by a high expansion of zone 1, while pigmentation was apparently not affected. Finally, the analysis of Δ*fis* mutant strain revealed a colony where the zone 1 (expansion zone) could not be detected. Additionally, this strain presented a darker zone 3, which could be indicative of higher production of curli fimbriae than wild-type strain, with addition of glucose to the growth media only generated small changes in the colony such as an apparent reduction in size.

Altogether, morphology assays have shown that IHF and Fis GRs are important contributors to proper biofilm development, as could be indicated by drastic changes in the three well-characterized zones in these mutants. In contrast, the role of CRP protein (which was indirectly assayed by the addition of glucose to the media) during mature biofilm formation could not be dependably inferred, since the change in the nutritional state of the colonies could have a strong impact in the process. Taken together, these data indicate that these GRs could have an important role in the final biofilm structure in *E. coli* that could be the result of the detected change in gene expression of key regulatory elements in the planktonic to biofilm regulatory network.

## Conclusions

The work presented here describes the systematic investigation of the regulatory interaction mediated by GRs during the transition from planktonic to biofilm in the bacterium *E. coli*. While several works have addressed this issue before, data generated has been fragmented and few targets have been investigated at each time. The systematic investigation used here allowed the identification of novel interactions mediated by CRP, Fis and IHF. More importantly, the modulation of some critical nodes, such as *fliA* by IHF, could explain the strong phenotypic effects observed for motility and attachment assays, as this regulator has also been found to be a modulator of genes involved in the biofilm program as *rpoS*, *matA* and *csgD* ^33, 38, 39, 40^. The molecular mechanism involved in the different phases of biofilm formation by IHF remains to be determined. However, in this work, we present evidence that IHF is a key element to biofilm formation by modulating gene expression level of the flagella-biofilm program, suggesting that IHF could be a good candidate to disrupt/engineer bacterial biofilm structure. Here, we also demonstrated that Fis plays a more major role in the biofilm formation network than anticipated, both using promoter assays as well as physiological tests. The evidence in this study indicates that, CRP, IHF and Fis are important effectors that modulate the flagella-biofilm network. Interestingly, no condition in this study shows absence of motility or biofilm formation, suggesting that, while these GRs are important modulators of these processes, they are not essential genes to suppress completely the flagella-biofilm program. Additionally, it has been demonstrated that bacteria without the main effectors *rpoS* or *csgD* (also *rpoE*, *csgB*, *flhDC* or *fliA*) are still capable of producing a biofilm ^7,41^. Altogether, the high plasticity observed in the flagella-biofilm program clearly shows that the network has evolved the robustness to compensate for the loss of critical nodes, indicating the importance of this program for bacterial survival. Finally, the systematic identification of novel connections for a specific network will allow prediction of the behavior of the biological system in the absence of different TFs. It opens the possibility of using genetic engineering to achieve a balanced connectivity between synthetic circuits and the entire bacterial network when performing a specific task. In other words, that rational interactive synthetic circuits can operate in accordance with the natural network of a specific organism, thereby allowing enhanced performance of a desired biological task.

## EXPERIMENTAL PROCEDURES

### Bacterial strains, media and growth conditions

The list of bacterial strains, plasmids and primers is presented in **Table 1**. *Escherichia coli* wild-type (BW25113) or strains genetically depleted of IHF or Fis GRs ^42^ were used as the host of all plasmids. *E. coli* DH5a or DH10b were used as cloning strains. DNA of *E. coli* BW25113 and MG1655 were used as templates. *E. coli* cells were grown in Luria Broth (LB) or M9 minimal media (6.4g/L Na_2_HPO_4_·7H_2_O, 1.5g/L KH_2_PO_4_, 0.25g/L NaCl, 0.5g/L NH_4_Cl) supplemented with 2mM MgSO_4_, 0.1mM CaCl_2_, 0.1mM casamino acids, and 1% glycerol or 0.4% glucose as the carbon source. Chloramphenicol was added (34 μg/mL) to ensure plasmid maintenance. Cells were grown at 37°C with constant shaking at 220 rpm overnight.

**Table 1.**
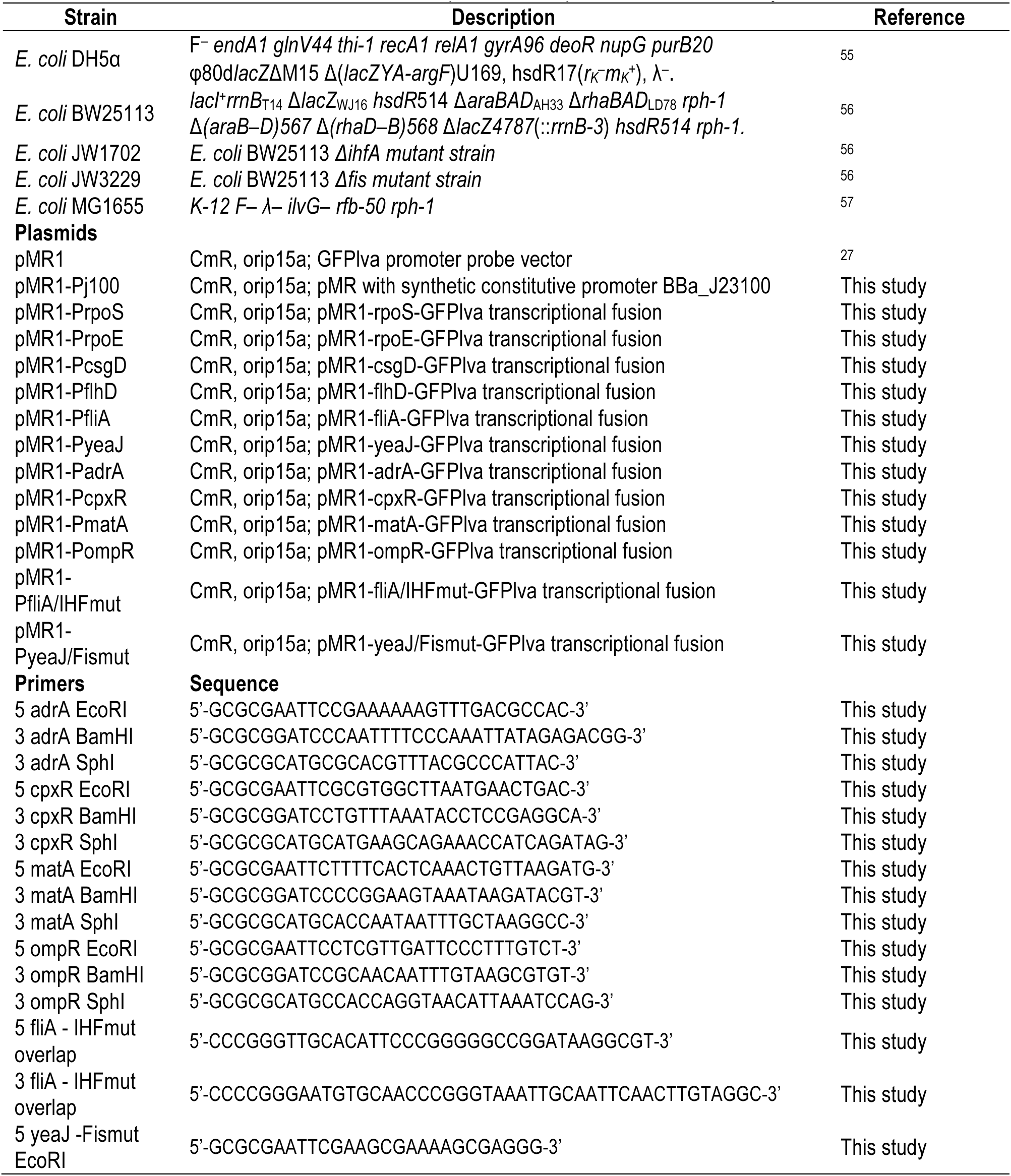
Bacterial strains, plasmids and primers used in this study.

### Plasmid construction

Plasmids and primers used in this work are listed on Table 2 and 3 respectively. As a control of all experiments we transformed *E. coli* BW25113 wild-type and mutant strains with an empty pMR1 vector ^27^ and pMR1-Pj100. To analyze the signal integration of GRs at the studied genes, we amplified the promoter regions by PCR and cloned into pMR1 vector using standard protocols ^43^. DNA from *E. coli* BW25113 wild-type was used as a template for all constructions except for *flhD* promoter, for which *E. coli* MG1655 was used. PCR was performed using 50 ng of DNA template with 50pmol of each primer (purchased from Sigma and extended) and 1U Phusion® High-Fidelity DNA Polymerase (New England Biolabs M0530L) in a 50 μl reaction volume. Initial denaturation was at 98°C for five minutes; subsequently, amplification was done over 30 cycles at 98°C, 60°C and 72°C, each with duration of 30 seconds. The final extension step was performed at 72°C for ten minute. All PCR products were gel purified and cloned as EcoRI/BamHI fragments in pMR1, previously digested with the same enzymes. The resulting plasmids were DNA sequence verified and correct plasmids were used to transform *E. coli* BW25113 (wild-type), *E. coli* JW1702 (*Δihf*) and *E. coli* JW3229 (*Δfis*).

### Promoter activity assay

For promoter activity measurements, single colonies of wild-type and mutant *E. coli* strains harboring the different plasmids were picked from fresh plates. Each strain was inoculated and grown overnight at 37 °C in 2 mL of M9 medium with glycerol or glucose containing chloramphenicol with shaking at 225 rpm. Stationary phase cultures were diluted to a final optical density (OD) of 0.05 in 200 μL of M9 medium containing either glycerol or glucose as required, and supplemented with chloramphenicol. Cell growth and GFP fluorescence was quantified every 20 minutes over an 8 hr incubation at 37 °C using Victor X3 plate reader (PerkinElmer, Waltham, Massachusetts, USA). Under these conditions, all strains presented similar growth rates (**Fig. S8**). Data were analyzed from at least three biological samples, arbitrary units were calculated by dividing GFP-corrected by OD600-corrected, subsequently, the mean and standard error for each promoter was calculated and graphed using Microsoft Excel and R software.

### Bacteria motility assay

The effect of GRs on motility was evaluated as following. *E. coli* wild-type or mutant strains harboring pMR1 plasmid were grown overnight at 37°C on LB plates. Inoculation of a single colony onto motility plates (tryptone 1%, NaCl 0.25%, agar 0.3% and when indicated glucose was added) was done by using a toothpick ^44^. The motility halos were measured at 18h and 24h. Each strain was evaluated in triplicate from independent plates.

### Biofilm formation in liquid media

Biofilm formation was assayed using the Microtiter Dish Biofilm Formation Assay ^45^. Single colonies of *E. coli* wild-type or mutant strains harboring all the constructions were grown overnight at 37°C in M9 supplemented with glycerol or glucose. The cultures were diluted 1:100 in 200μl of fresh M9 media with either glycerol or glucose. A 96-well plate was incubated at 37°C for 72h, after which cells were gently washed four times with distilled water and left at room temperature for 15 minutes with 200μL of crystal violet 0.1% w/v solution. Subsequently, vigorous washing was performed with distilled water (four times) and the plates left to dry at room temperature for 24h. Finally, 200μL acetic acid 30% w/v was added and after 15 minutes, the solution was transferred to a new plate and Optical Density at 550nm was measured in FLUOstar Omega (BMG Labtech) plate reader. All the culture dilutions, staining and quantification steps were performed by using procedures developed for a robotic platform called Edwin ^46^. Data from biological triplicates were analyzed by Microsoft Excel software. Average values and standard deviation was determined.

### Biofilm formation in solid media

Biofilm morphology was evaluated by Congo red assay ^47^. Chloramphenicol was added during all steps to ensure plasmid maintenance. Wild-type and mutant *E. coli* strains with pMR1 plasmid were grown overnight at 37°C on LB agar plates. Single colonies were picked and grown for 14h at 37°C on 1ml LB at 220rpm. The cultures were washed twice with MgSO4 and resuspended in M9 media with either glycerol or glucose as required, to an OD at 600nm (~ 0.5). 5 μL drop of each culture was added into YESCA-CR plates ^48^. YESCA media (1 g/L yeast extract, 20 g/L agar) was complemented with Congo red 50 μg/ml diluted in KPi buffer (50 mM potassium phosphate buffer, pH 7.2, 28.9 mM KH2PO4, 21.1 mM K2HPO4 in water), where indicated glucose was added at 0.4% final concentration. YESCA-CR seeded plates were allowed to dry in a sterile environment and, left to grow at 37°C. Biofilm development was followed (documented) for 6 days using a fluorescence stereoscope (Leica MZ16). Biological triplicate images were analyzed by ImageJ and prisma software was used to determine the statistical significance of the experiments by using one-way ANOVA.

### Network Construction

To generate the transcriptional regulatory network that will define our genes of interest, we gathered information from research articles that contained information on genes that modulate the transition between planktonic to biofilm formation. Eventually, the data were compared with information available at Regulon DB, EcoCyc DB and Kegg pathway DB. Cytoscape 3.4.1 was used to generate the transcriptional network with slight modifications ^49^. Briefly, we transformed our mined data into activation or inhibition statements for each of the partner molecules. Secondly, to construct the network graph, we used Cytoscape 3.4.1 in which different connections were added to each node on the collected data. Next, networks were analyzed using different algorithms, such as degree, out-degree, betweenness centrality by using Cytoscape 3.4.1 software. Finally, we transformed our GFP dynamics gene expression experimental results into functional interactions (inhibits or activates). Results with same function as reported were maintained without changes in the rewired network, while those with different activity were changed in the network. During the data collection from the literature, some TFs were reported with dual effect (activation and repression) for the same target gene. For instance, *rpoS* expression was reported to be activated but also inactivated by the CRP global regulator^50, 51, 52^. Additionally, most of the genes that were reported as being regulated by different TFs, but in the web-databases the same gene did not present a *cis*-element to any TFs. To overcome those constrains we considered for the network that any TF, master regulator, global regulator, small molecule or RNAs could be a node. This last, despite being gathered, were not included because RNA-node analysis of regulatory gene expression significantly increases the complexity of the network ^12,53^ and because of the complex requirements of molecular tools to evaluate this topic. For edges direction analysis, we considered the activation or inhibition most reported, or both when divergent reports were found. The data were plotted in a power-law graph using an organic algorithm to give cluster visualization generating the transcriptional regulatory network of both planktonic and biofilm structure. Furthermore, we performed degree analysis in order to present a general view of the most connected nodes, followed by edge and node betweenness analysis ^54^ in order to disclose the main effectors and the logic pathway that could describe the network. Finally, we also performed the out degree analysis at the integrated network to identify the potential talkative nodes.

## ACKNOWLEDGMENTS

The authors express their thanks to members of the Silva-Rocha & Elfick labs for insightful discussion and suggestions in developing this work. This work was supported by a FAPESP Young Research Award (grant number 2012/22921-8) and by a Royal Society Newton International Exchange award (NI140137). GRA was supported by a CAPES PhD fellowship while ASM was supported by FAPESP Scientific Initiation Award (2015/22386-3). AdlH was supported by an EPSRC Programme Grant (EP/J02175X) to the Flowers Consortium.

## AUTHOR CONTRIBUTIONS

G.R.A. and R.S.R. conceived the work. G.R.A., A.dH. and A.S.M. performed the experiments. G.R.A., A.dH. and A.M.S. analyzed the data. G.R.A. and R.S.R. drafted the manuscript. A.dH. and A.E. corrected the manuscript. A.E. and R.S.R. provided funding for the execution of the work. All authors read and approved the manuscript.

## MATERIAL & CORRESPONDENCE

Correspondence and material request should be addressed to Rafael Silva Rocha.

## ADDITIONAL INFORMATION

Supplementary Information accompanies this paper online.

